# Chronic alcohol abuse affects the clinical course and outcome of community-acquired bacterial meningitis

**DOI:** 10.1101/567347

**Authors:** Marcin Paciorek, Agnieszka Bednarska, Dominika Krogulec, Michał Makowiecki, Justyna D Kowalska, Dominik Bursa, Anna Świderska, Joanna Puła, Joanna Raczyńska, Agata Skrzat-Klapaczyńska, Marek Radkowski, Tomasz Laskus, Andrzej Horban

## Abstract

**Background:** The aim of the study was to determine the effect of chronic alcohol abuse on the course and outcome of bacterial meningitis (BM).

**Materials/methods:** We analyzed records of patients with BM who were hospitalized between January 2010 and December 2017 in the largest neuroinfection center in Poland.

**Results:** 340 adult patients (211 men and 129 women) were analyzed. Forty-five (13.2%) patients were alcoholics (39 men and 6 women). Compared to non-alcoholics, alcoholics were more likely to present with seizures (33.3% vs 12.6%, p<0.001), scored higher on the Sequential Organ Failure Assessment (SOFA) (median 3 vs 2, p<0.001) and lower on the Glasgow Coma Scale (GCS) (median 10 vs 12, p<0.001) and had worse outcome as measured by the Glasgow Outcome Score (GOS) (median 3 vs 5, p<0.001). Furthermore, alcoholics were less likely to complain of headache (23.3% vs 52.3%, p<0.001) and nausea/vomiting (11.4% vs 33.6%, p=0.005) and had lower concentration of glucose in cerebrospinal fluid (CSF) (median 0,58 mmol/L vs 1,97, p=0.025). In the multiple logistic regression analysis, alcoholism was independently associated with lower GCS (OR 0.716, 95% CI 0.523-0.980, p=0.036), presence of seizures (OR 4.580, 95% CI 1.065-19.706, p=0.041), male gender (OR 4.617, 95% CI 1.060-20.113, p=0.042) and absence of nausea/vomiting (OR 0.205, 95%CI 0.045-0.930, p=0.040). Furthermore, alcoholism (regression coefficient [−0.636], 95% CI [− 1.21] – [−0.06], p=0.031), lower GCS score (regression coefficient 0.144, 95% CI 0.06-0.23, p=0.001) and higher urea blood concentration (regression coefficient [−0.052], 95% CI [−0.10] – [−0.01], p=0.018) were independently associated with worse outcome measured by GOS.

**Conclusions:** Compared to non-alcoholics, chronic alcohol abusers are more likely to present with seizures, altered mental status, higher SOFA score and have an increased risk of unfavorable outcome. In multivariate analysis seizures and low GCS were independently associated with alcoholism, while alcoholism was independently associated with worse outcome.

## Introduction

Bacterial meningitis is a life-threatening medical emergency characterized by high mortality rate and frequent long-term neurological sequelae ^(1)^; in Poland, where the registration of BM cases is mandatory, the annual incidence of BM in the years 2010-2017 ranged between 1.97/100,000 and 2.5/100,000 ^(2–4)^ which is higher than is some other European countries such as Finland (0.7/100,000) Netherlands (0.94/100,000) or England and Whales (1.44/100,000) ^(5–7)^. In Poland alcohol per capita consumption (in liters of pure alcohol) for population ≥ 15 years old was 11.4 in 2010 and 11.6 in 2016 which is similar to many other European countries ^(8)^. Chronic alcohol abuse may have adverse medical consequences such as stroke, cancers, liver cirrhosis, gastrointestinal disorders, cognitive deficits, and peripheral neuropathy ^(9)^. In addition, alcoholics are more susceptible to bacterial infections and these infections carry a worse prognosis. There is evidence that the relative risk of bacterial pneumonia correlates with the level of alcohol intake ^(10)^. While this may be in part a consequence of lifestyle and malnutrition, there is ample evidence that chronic alcohol abuse itself may impair various host immune responses; for example it was demonstrated to negatively affect phagocytosis and superoxide production in alveolar macrophages after bacterial challenge and leads also to hypo-responsiveness of neutrophils to chemotactic signals ^(11, 12)^. The initial diagnosis of BM is typically based on clinical signs and symptoms and antibiotic treatment is usually initiated before the etiological factor is identified as prompt therapy is crucial for improving the outcome ^(13, 14)^.

The aim of our study was to determine whether chronic alcohol abuse has any impact on clinical manifestations, etiologic factors and outcome in BM. Such an analysis is rare in the literature and was confined so far to two studies from Netherlands ^(15, 16)^. We found that alcoholics may have a different clinical presentation and worse outcome compared to non alcoholic patients.

## Materials and Methods

We evaluated records of adult (≥ 18 years old) patients with community-acquired BM who were admitted to the Hospital for Infectious Diseases in Warsaw from January 1st, 2010 to December 31st, 2017. Only patients who underwent a diagnostic lumbar puncture were included in the analysis. The diagnosis of BM was based on fulfilling at least one of the following criteria: positive cerebrospinal fluid (CSF) culture, positive CSF Gram staining, typical CSF findings (pleocytosis ≥ 100 cells/μl with≥ 90% neutrophils and decrease of CSF glucose level < 2.2 mmol/L). Patients with CSF findings typical for BM but negative blood and CSF culture and negative microscopic CSF examination were considered to have BM of unknown etiology. Patients with positive blood culture but negative CSF culture and negative CSF microscopic examination were considered to have BM caused by pathogen cultured from blood.

Initial antimicrobial treatment followed current guidelines ^(17)^ and included vancomycin and a third generation cephalosporin in patients below 50 years old and a combination of ampicillin and vancomycin together with a third generation cephalosporin in those ≥50 years or those likely to be immunocompromised (patients with cancer, diabetes mellitus, HIV infection, liver cirrhosis, alcoholics, patients receiving immunosuppressive therapy). Diagnosis of tuberculous meningitis was based on at least one of the following: positive culture, positive nucleic acid amplification, positive Ehrlich-Ziehl-Neelsen staining of CSF. Patients with tuberculous meningitis were treated with rifampicin, isoniazid, pyrazinamide and ethambutol or streptomycin. All patients received adjunctive therapy with dexamethasone. Patients with meningitis secondary to head trauma, neurosurgical procedures as well as hospital-acquired infections were excluded from analysis.

Alcoholism was diagnosed according to the WHO criteria ^(18)^ as a chronic continual or periodic consumption of alcohol characterized by impaired control over drinking, frequent episodes of intoxication, and preoccupation with alcohol and the use of alcohol despite adverse consequences. Questions about alcohol consumption were a standard part of medical interview, but there were no questions relating to the quantity of alcohol intake per day or week.

The U Mann Whitney test was used to compare continuous variables and the Chi-square test was used to evaluate nominal variables. A p value of <0.05 was considered significant. Logistic regression was used to calculate adjusted odds ratios and to determine variables independently associated with alcoholism and those independently influencing outcome reflected by Glasgow Outcome Scale (GOS). Since GOS is not a nominal but continuous variable, we used general linear model to calculate the coefficients for this analysis. Statistical analysis were performed using program R version 3.5.2 ^(19)^.

## Results

The final analysis included 340 patients with bacterial meningitis (211 men and 129 women, median age 57, interquartile range [IQR] 41-69). Among this group 45 (13.2%) patients were considered alcoholics (39 men and 6 women, median age 53, IQR 44-59). The proportion of men was significantly higher among alcoholic patients than among non-alcoholics (86% vs 42%, p<0.001).

At admission, alcoholic patients were more likely to present with seizures (33.3% vs 12.6, p<0.001) (Table 1) but were less likely to complain of headache (23.3% vs 52.3%, p<0.001) and nausea/vomiting (11.4% vs 33.6%, p=0.005). Furthermore, compared to non-alcoholic patients, they were in worse clinical condition: they scored higher on the Sequential Organ Failure Assessment (SOFA) (median 3 [IQR 2-6] vs median 2 [IQR 1-5], p<0.001) and lower on the Glasgow Coma Scale (GCS) (median 10 [IQR 7-12] vs median 12 [IQR 9-14], p<0.001). Alcoholic patients were also more likely to require Intensive Care Unit (ICU) admission (48.9% vs 37.6 %) and their mortality was higher (24.4% vs 15.4%), but these differences did not reach statistical significance. The main reasons for ICU admission were: respiratory failure, coma, and septic shock. The clinical outcome reflected by the GOS was significantly worse among alcoholics (median 3 [IQR 1-5] vs median 5 [IQR 3-5]; p < 0.001; Table 1).

**Table 1.**
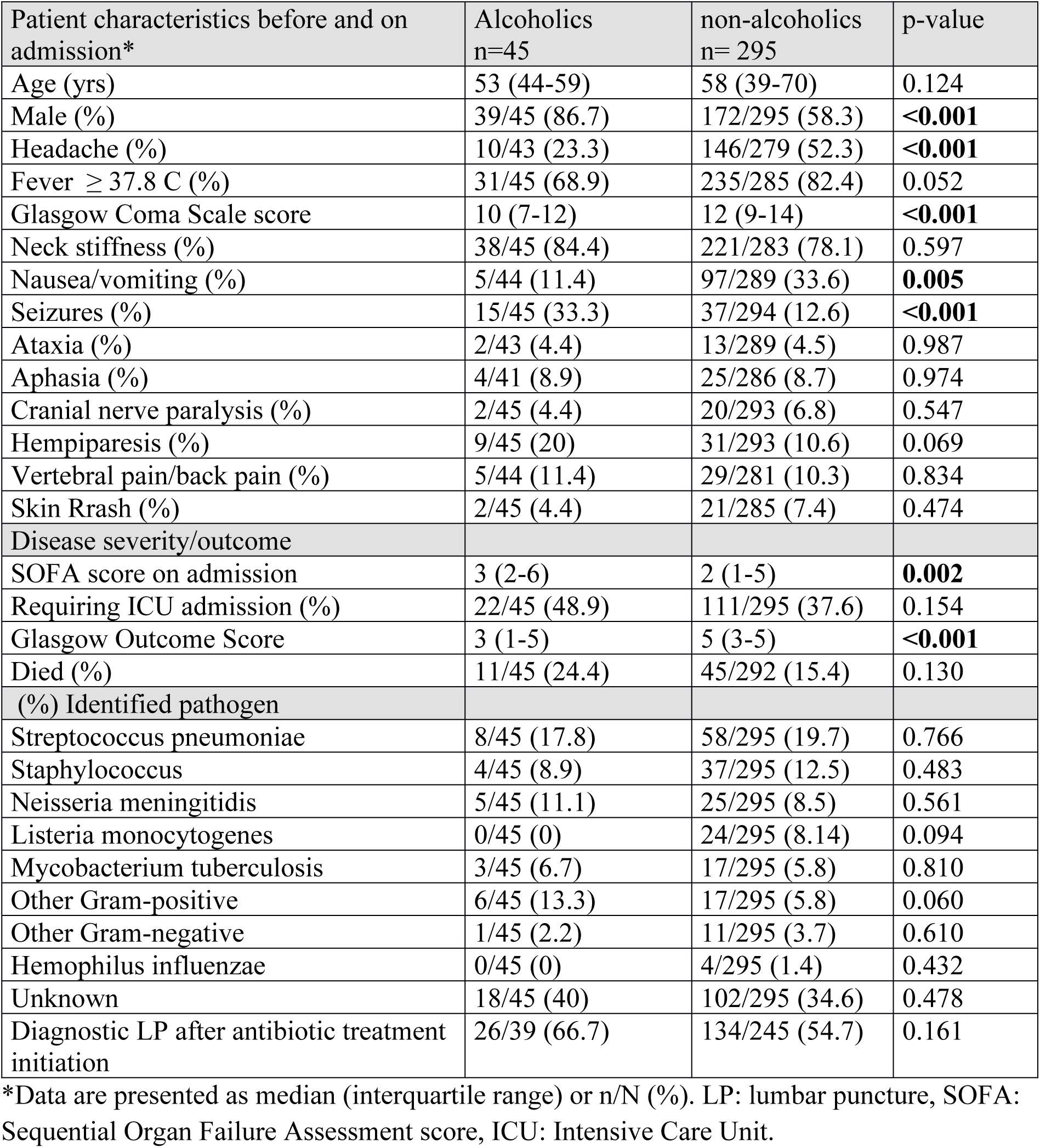
Demographic, clinical and etiological data in alcoholic and non–alcoholic patients with bacterial meningitis.

Analysis of laboratory parameters (Table 2) revealed that alcoholic patients had significantly higher serum concentration of the D-dimers (median 3273 μl/L [IQR 1751-5817] vs median 2241 μl/L [IQR 1159-4262], (p=0.013) and lower concentration of glucose in CSF (median 0.58 mmol/L [IQR 0-2.3] vs median 1.97 [IQR 0.11-3.40], p=0.025; Table 2).

**Table 2.**
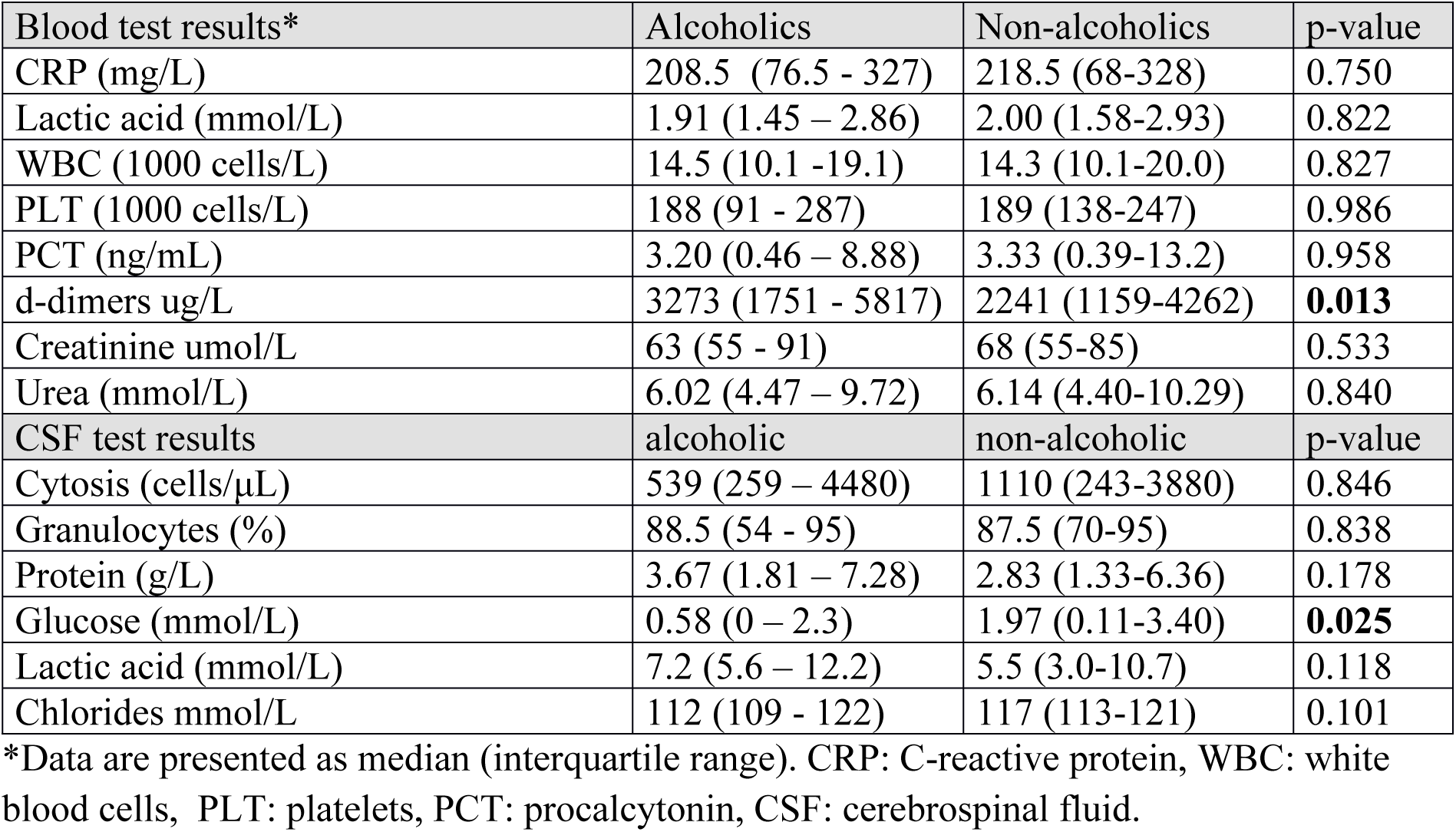
Laboratory blood and cerebrospinal fluid (CSF) results in alcoholic and non-alcoholic patients with bacterial meningitis.

In multiple logistic regression analysis (Table 3) alcoholism was independently associated with a lower GCS score (OR 0.716, 95% CI 0.523-0.980, p=0.036), male gender (OR 4.617, 95% CI 1.060-20.113, p=0.042), the presence of seizures (OR 4.580, 95% CI 1.065-19.706, p=0.041) and absence of nausea/vomiting (OR 0.205, 95% CI 0.045-0.930, p=0.040).

**Table 3.** Multiple logistic regression analysis of factors independently associated with alcoholism in patients with bacterial meningitis.

| Variable* | P value | OR | 95% CI |
| --- | --- | --- | --- |
| GCS | **0.036** | 0.716 | 0.523-0.980 |
| GOS | 0.780 | 0.933 | 0.575-1.515 |
| SOFA | 0.075 | 0.814 | 0.649-1.021 |
| CSF glucose | 0.335 | 0.850 | 0.611-1.183 |
| Male gender | **0.042** | 4.617 | 1.060-20.113 |
| Headache | 0.316 | 0.535 | 0.157-1.819 |
| Nausea/vomiting | **0.040** | 0.205 | 0.045-0.930 |
| Seizures | **0.041** | 4.580 | 1.065-19.706 |

\*Results are presented as odds ratio (OR) and confidence interval (CI).
GCS: Glasgow Coma Scale, GOS: Glasgow Outcome Scale, SOFA: Sepsis related Organ Failure Assessment score, CSF glucose: concentration of glucose in cerebrospinal fluid.

Furthermore, alcoholism, lower GCS score and higher urea blood concentration were independently associated with worse prognosis as assessed by GOS (Table 4).

**Table 4.** Multiple logistic regression analysis of factors independently associated with lower Glasgow Outcome Score (GOS) in patients with bacterial meningitis.

| Variable | Regression coefficient | 95% CI | P value |
| --- | --- | --- | --- |
| Alcoholism | -0.636 | (-1.209) – (-0.063) | **0.031** |
| Seizures | 0.180 | (-0.037) – 0.728 | 0.521 |
| Etiology unknown | 0.100 | (-0.353) – 0.552 | 0.667 |
| Str. pneumoniae | -0.010 | (-0.669) – 0.470 | 0.732 |
| N. meningitidis | 0.512 | (-0.244) – 1.267 | 0.186 |
| Age | -0.009 | (-0.021) – 0.003 | 0.161 |
| GCS | 0.144 | 0.061 – 0.226 | **0.001** |
| SOFA | -0.068 | (-0.160) – 0.025 | 0.154 |
| CRP | 0.001 | (-0.001) – 0.002 | 0.841 |
| PLT | 0.001 | (-0.001) – 0.002 | 0.633 |
| PCT | -0.001 | (-0.008) – 0.005 | 0.665 |
| Urea | -0.052 | (-0.095) – (-0.009) | **0.018** |
| CSF protein | -0.007 | (-0.040) – 0.025 | 0.656 |
| CSF glucose | 0.034 | (-0.061) – 0.129 | 0.484 |

CI: confidence interval, Str. Pneumoniae: Streptococcus pneumoniae, N. meningitidis: Neisseria meningitidis, GCS: Glasgow Coma Scale, SOFA: Sepsis related Organ Failure Assessment score, PLT: platelet level, PCT: concentration of procalcitonin in blood, CSF protein: concentration of protein in cerebrospinal fluid, CSF glucose: concentration of glucose in cerebrospinal fluid.

An etiological factor was identified only in 60% of alcoholics and 35% of non-alcoholics (Table 1). However, in 26 out of 39 alcoholics (67%) and in 134 out of 245 non-alcoholics (55%) antibiotic treatment initiation preceded diagnostic spinal tap. Streptococcus pneumoniae was the most common causative organism in both groups followed by Staphylococci, Neisseria meningitidis, and Mycobacterium tuberculosis. Listeria monocytogenes was identified only in non-alcoholic patients but this difference did not reach statistical significance (Table 1).

## Discussion

According to national survey data ^(20)^, alcoholics constitute approximately 2% of the Polish general adult population, but among our patients with BM this proportion was as high as 13% However, since self-reporting survey methods may lead to underestimation of drinking levels this proportion could be even higher. This high prevalence of alcoholics among our patients might be a direct consequence of impaired immune response to bacterial pathogens related to chronic alcohol abuse ^(21–24)^. Thus, it is likely that alcoholism is a risk factor for the development of BM, much like for the bacterial pneumonia. In the nationwide cohort study conducted in the Netherlands the proportion of alcoholics in BM patients was 6% which was also higher than in general population ^(8, 15^

Compared to non-alcoholics, alcohol abusers in our study were less likely to present with high fever, headache, and nausea. The absence of such typical symptoms of BM ^(25)^ might cause diagnostic problems and in effect result in the delay of treatment initiation, thus worsening patients’ outcome prospects. Interestingly, attenuated fever response to lipopolysaccharide and interleukin 1β has been described experimentally in rats after they were fed alcohol for a prolonged period ^(26)^, while lower incidence of headache among alcoholics with BM seems to be a common phenomenon as it was previously reported by van Veen et al ^(15)^ in their population-based cohort study.

Another and most striking difference in the clinical presentation was significantly higher incidence of seizures among alcoholic patients (33% vs 13%; p < 0.001). While high incidence of seizures among alcohol abusers presenting with BM is a well known phenomenon, the numbers in previous reports were lower and ranged from 13% to 18% ^(15, 16)^. Furthermore, in our multivariate analysis the presence of seizures was independently associated with alcoholism, In a large prospective cohort study of community-acquired BM in Netherlands, seizures were associated with higher mortality rate ^(27)^. The reasons for the latter are not entirely clear, but seizures in BM patients could reflect more severe inflammatory damage ^(28, 29)^ or alcohol withdrawal syndrome, which in itself carries increased mortality risk^(30)^. However, in our study seizures were not independently associated with outcome.

The presence of seizures and other signs and symptoms associated with the alcohol withdrawal syndrome or even alcohol intoxication itself could not only complicate clinical presentation, but could also negatively affect consciousness level: in our study alcoholic patients scored significantly lower on the GCS scale at admission compared to non-alcoholics. These findings are compatible with previous studies of van Veen et al ^(15)^ and Weisfelt et al ^(16)^ who found higher prevalence of coma, defined as GCS below 8 points, and lower GCS score, respectively, among alcoholic patients with BM when compared to non-alcoholics. However, the above differences did not reach statistical significance. Obviously, more pronounced impairment in consciousness among alcoholics could be also related to more severe inflammation process.

Alcoholic patients in our study scored higher on SOFA at admission. SOFA scoring is currently a crucial part of sepsis definition ^(31)^ as it reflects the level of organ dysfunction and correlates with patients’ outcome. However, its use is not confined to sepsis as in two previous studies a high SOFA score combined with low GCS ^(32, 33)^ were found to be predictors of unfavorable outcome in BM. The mortality rate among alcoholic patients in our analysis was high (24%), and exceeded the death rate in non-alcoholic patients (15%), although this difference did not reach statistical significance (p=0.13). While alcohol abuse was not an independent predictor for mortality in multivariate analysis, it was associated with worse outcome, as measured by the GOS. These results are similar to those reported by Wiesfelt et al ^(26)^ who also found in multivariate analysis that alcoholism is associated with unfavorable outcome but not with higher mortality. The reasons for the worse outcome among alcoholics are probably manifold including coexisting alcohol withdrawal syndrome ^(34)^, alcohol-related encephalopathy, malnutrition and the presence of seizures. Furthermore, analogous to bacterial pneumonia, the course of BM may simply be more severe due to profound negative effects of chronic alcohol use on the efficiency of immune response ^(35)^.

In addition to alcohol, also low GCS score and high blood urea levels were independently associated with worse outcome (Table 4). The influence of GCS on outcome is not surprising and was previously reported by others ^(32, 33)^, while increased urea might identify a subset of patients with multiple organ failure. Notably, SOFA score was also predictive of worse outcome, although it did not reach statistical significance. With the exception of blood D-dimer levels and CSF glucose no differences were found in the results of routine tests between alcoholic and non-alcoholic patients. High blood D-dimer levels are common in chronic alcohol abusers and could be the effect of hemostatic system activation related to oxidative stress ^(36)^. Interestingly, low concentration of glucose in the CSF was previously correlated with adverse clinical outcomes in patients with BM ^(37)^.

Streptococcus pneumoniae was the most common causative microorganism (19.4%) in our patients (Table 1) but its prevalence was not as high as in some other European studies ^(38–40)^. Such relatively low prevalence of Streptococcus pneumoniae is not a local phenomenon confined to our center, as it accounted for only 22% of BM cases registered in Poland ^(3)^. Interestingly, similar prevalence of Streptococcus pneumoniae and Staphylococci was reported by Ishihara et al.^(41)^ in the Japanese population of BM patients. However, similar to our study, in the majority of patients antibiotic treatment was initiated before CSF collection. It should be pointed out that nucleic acid amplification assays were not used for pathogen identification in our study, with the exception of cases suspected of tuberculous meningitis. Such techniques might be especially useful for the identification of pathogens in patients in whom empirical antibiotic therapy preceded collection of relevant biological samples. Unexpectedly, Listeria monocytogenes was not found to in any alcoholic patients in our study, although it was a common pathogen constituting 7% of all cases in the non-alcoholic group. This results are in contrast to other studies, in which alcoholism was identified as one of the predisposing factors for listeriosis ^(25, 42)^. The reasons for this are unclear.

In conclusion, we found that alcoholic patients with BM, when compared to their non-alcoholic counterparts, are more likely to present with seizures and more severely altered mental status and have an increased risk for unfavorable outcome.

## References

1. van de Beek D, Brouwer MC, Thwaites GE, Tunkel AR. Advances in treatment of bacterial meningitis. Lancet. 2012;380(9854):1693–702.

2. Paradowska-Stankiewicz I, Piotrowska A. Meningitis and encephalitis in Poland in 2014. Przegl Epidemiol. 2016;70(3):349–57.

3. Paradowska-Stankiewicz I, Piotrowska A. Meningitis and encephalitis in Poland in 2015. Przegl Epidemiol. 2017;71(4):493–500.

4. http://wwwold.pzh.gov.pl/oldpage/epimeld/2017/Ch_2017_wstepne_dane.pdf.

5. Polkowska A, Toropainen M, Ollgren J, Lyytikainen O, Nuorti JP. Bacterial meningitis in Finland, 1995-2014: a population-based observational study. BMJ Open. 2017;7(5):e015080.

6. Bijlsma MW, Brouwer MC, Kasanmoentalib ES, Kloek AT, Lucas MJ, Tanck MW, et al. Community-acquired bacterial meningitis in adults in the Netherlands, 2006-14: a prospective cohort study. Lancet Infect Dis. 2016;16(3):339–47.

7. Okike IO, Ribeiro S, Ramsay ME, Heath PT, Sharland M, Ladhani SN. Trends in bacterial, mycobacterial, and fungal meningitis in England and Wales 2004-11: an observational study. Lancet Infect Dis. 2014;14(4):301–7.

8. https://apps.who.int/iris/bitstream/handle/10665/274603/9789241565639-eng.pdf

9. Schuckit MA. Alcohol-use disorders. Lancet. 2009;373(9662):492–501.

10. Samokhvalov AV, Irving HM, Rehm J. Alcohol consumption as a risk factor for pneumonia: a systematic review and meta-analysis. Epidemiol Infect. 2010;138(12):1789–95.

11. Brown LA, Harris FL, Ping XD, Gauthier TW. Chronic ethanol ingestion and the risk of acute lung injury: a role for glutathione availability? Alcohol. 2004;33(3):191–7.

12. Sachs CW, Christensen RH, Pratt PC, Lynn WS. Neutrophil elastase activity and superoxide production are diminished in neutrophils of alcoholics. Am Rev Respir Dis. 1990;141(5 Pt 1):1249–55.

13. Brouwer MC, Thwaites GE, Tunkel AR, van de Beek D. Dilemmas in the diagnosis of acute community-acquired bacterial meningitis. Lancet. 2012;380(9854):1684–92.

14. Domingo P, Pomar V, Benito N, Coll P. The changing pattern of bacterial meningitis in adult patients at a large tertiary university hospital in Barcelona, Spain (1982-2010). J Infect. 2013;66(2):147–54.

15. van Veen KE, Brouwer MC, van der Ende A, van de Beek D. Bacterial meningitis in alcoholic patients: A population-based prospective study. J Infect. 2017;74(4):352–7.

16. Weisfelt M, de Gans J, van der Ende A, van de Beek D. Community-acquired bacterial meningitis in alcoholic patients. PLoS One. 2010;5(2):e9102.

17. koroun.edu.pl/wp-content/uploads/2017/10/Rekomendacje-ukl-nerwowy_2011.pdf

18. http://www.who.int/substance_abuse/terminology/who_lexicon/en/.

19. https://www.r-project.org.

20. http://www.parpa.pl/index.php/33-analizy-badania-raporty/132-statystyki.

21. Bhatty M, Pruett SB, Swiatlo E, Nanduri B. Alcohol abuse and Streptococcus pneumoniae infections: consideration of virulence factors and impaired immune responses. Alcohol. 2011;45(6):523–39.

22. Jerrells TR, Slukvin I, Sibley D, Fuseler J. Increased susceptibility of experimental animals to infectious organisms as a consequence of ethanol consumption. Alcohol Alcohol Suppl. 1994;2:425–30.

23. Gandhi JA, Ekhar VV, Asplund MB, Abdulkareem AF, Ahmadi M, Coelho C, et al. Alcohol enhances Acinetobacter baumannii-associated pneumonia and systemic dissemination by impairing neutrophil antimicrobial activity in a murine model of infection. PLoS One. 2014;9(4):e95707.

24. Karavitis J, Kovacs EJ. Macrophage phagocytosis: effects of environmental pollutants, alcohol, cigarette smoke, and other external factors. J Leukoc Biol. 2011;90(6):1065–78.

25. van de Beek D, de Gans J, Spanjaard L, Weisfelt M, Reitsma JB, Vermeulen M. Clinical Features and Prognostic Factors in Adults with Bacterial Meningitis. New England Journal of Medicine. 2004;351(18):1849–59.

26. Taylor AN, Tio DL, Heng NS, Yirmiya R. Alcohol consumption attenuates febrile responses to lipopolysaccharide and interleukin-1 beta in male rats. Alcohol Clin Exp Res. 2002;26(1):44–52.

27. Zoons E, Weisfelt M, de Gans J, Spanjaard L, Koelman JH, Reitsma JB, et al. Seizures in adults with bacterial meningitis. Neurology. 2008;70(22 Pt 2):2109–15.

28. Lippai D, Bala S, Csak T, Kurt-Jones EA, Szabo G. Chronic alcohol-induced microRNA-155 contributes to neuroinflammation in a TLR4-dependent manner in mice. PLoS One. 2013;8(8):e70945.

29. Lippai D, Bala S, Petrasek J, Csak T, Levin I, Kurt-Jones EA, et al. Alcohol-induced IL-1beta in the brain is mediated by NLRP3/ASC inflammasome activation that amplifies neuroinflammation. J Leukoc Biol. 2013;94(1):171–82.

30. Rogawski MA. Update on the neurobiology of alcohol withdrawal seizures. Epilepsy Curr. 2005;5(6):225–30.

31. Singer M, Deutschman CS, Seymour CW, Shankar-Hari M, Annane D, Bauer M, et al. The Third International Consensus Definitions for Sepsis and Septic Shock (Sepsis-3). Jama. 2016;315(8):801–10.

32. Jordan I, Calzada Y, Monfort L, Vila-Perez D, Felipe A, Ortiz J, et al. Clinical, biochemical and microbiological factors associated with the prognosis of pneumococcal meningitis in children. Enferm Infecc Microbiol Clin. 2016;34(2):101–7.

33. Pietraszek-Grzywaczewska I, Bernas S, Lojko P, Piechota A, Piechota M. Predictive value of the APACHE II, SAPS II, SOFA and GCS scoring systems in patients with severe purulent bacterial meningitis. Anaesthesiol Intensive Ther. 2016;48(3):175–9.

34. Mirijello A, D’Angelo C, Ferrulli A, Vassallo G, Antonelli M, Caputo F, et al. Identification and management of alcohol withdrawal syndrome. Drugs. 2015;75(4):353–65.

35. Szabo G, Saha B. Alcohol’s Effect on Host Defense. Alcohol Res. 2015;37(2):159–70.

36. Trotti R, Carratelli M, Barbieri M, Micieli G, Bosone D, Rondanelli M, et al. Oxidative stress and a thrombophilic condition in alcoholics without severe liver disease. Haematologica. 2001;86(1):85–91.

37. Shrikanth V, Salazar L, Khoury N, Wootton S, Hasbun R. Hypoglycorrhachia in adults with community-acquired meningitis: etiologies and prognostic significance. Int J Infect Dis. 2015;39:39–43.

38. NR. Netherlands Reference Laboratory for Bacterial Meningitis (AMC/ RIVM). Bacterial meningitis in the Netherlands annual report 2010. Amsterdam: University of Amsterdam; 2011 [

39. Bodilsen J, Dalager-Pedersen M, Schonheyder HC, Nielsen H. Dexamethasone treatment and prognostic factors in community-acquired bacterial meningitis: a Danish retrospective population-based cohort study. Scand J Infect Dis. 2014;46(6):418–25.

40. Gjini AB, Stuart JM, Lawlor DA, Cartwright KA, Christensen H, Ramsay M, et al. Changing epidemiology of bacterial meningitis among adults in England and Wales 1991-2002. Epidemiol Infect. 2006;134(3):567–9.

41. Ishihara M, Kamei S, Taira N, Morita A, Miki K, Naganuma T, et al. Hospital-based study of the prognostic factors in adult patients with acute community-acquired bacterial meningitis in Tokyo, Japan. Intern Med. 2009;48(5):295–300.

42. Mook P, O’Brien SJ, Gillespie IA. Concurrent conditions and human listeriosis, England, 1999-2009. Emerg Infect Dis. 2011;17(1):38–43.

